# Gene–Dose–Dependent Reduction *Fshr* Expression Improves Spatial Memory Deficits in Alzheimer’s Mice

**DOI:** 10.1101/2024.02.16.580761

**Authors:** Funda Korkmaz, Steven Sims, Fazilet Sen, Farhath Sultana, Victoria Laurencin, Liam Cullen, Anusha Pallapati, Avi Liu, Satish Rojekar, George Penev, Ulliana Cheliadinova, Darya Vasilyeva, Guzel Burganova, Anne Macdonald, Mansi Saxena, Ki Goosens, Clifford Rosen, Orly Barak, Daria Lizneva, Anisa Gumerova, Keqiang Ye, Vitaly Ryu, Tony Yuen, Tal Frolinger, Mone Zaidi

**Author notes:** Corresponding Authors: Tal Frolinger; Tony Yuen and Mone Zaidi. Joint first authors (Funda Korkmaz, Steven Sims). Joint senior authors (Tony Yuen, Tal Frolinger, Mone Zaidi).

## Abstract

Alzheimer’s disease (AD) is a major progressive neurodegenerative disorder of the aging population. High post–menopausal levels of the pituitary gonadotropin follicle–stimulating hormone (FSH) are strongly associated with the onset of AD, and we have shown recently that FSH directly activates the hippocampal *Fshr* to drive AD–like pathology and memory loss in mice. To establish a role for FSH in memory loss, we used female *3xTg;Fshr*^+/+^*, 3xTg;Fshr*^+/–^ and *3xTg;Fshr^-/-^* mice that were either left unoperated or underwent sham surgery or ovariectomy at 8 weeks of age. Unoperated and sham–operated *3xTg;Fshr^-/-^*mice were implanted with 17β-estradiol pellets to normalize estradiol levels. Morris Water Maze and Novel Object Recognition behavioral tests were performed to study deficits in spatial and recognition memory, respectively, and to examine the effects of *Fshr* depletion. *3xTg;Fshr^+/+^*mice displayed impaired spatial memory at 5 months of age; both the acquisition and retrieval of the memory were ameliorated in *3xTg;Fshr^-/-^* mice and, to a lesser extent, in *3xTg;Fshr^+/-^* mice–– thus documenting a clear gene–dose–dependent prevention of hippocampal–dependent spatial memory impairment. At 5 and 10 months, sham–operated *3xTg;Fshr^-/-^* mice showed better memory performance during the acquisition and/or retrieval phases, suggesting that *Fshr* deletion prevented the progression of spatial memory deficits with age. However, this prevention was not seen when mice were ovariectomized, except in the 10–month–old *3xTg;Fshr^-/-^*mice. In the Novel Object Recognition test performed at 10 months, all groups of mice, except ovariectomized *3xTg;Fshr^-/-^* mice showed a loss of recognition memory. Consistent with the neurobehavioral data, there was a gene–dose–dependent reduction mainly in the amyloid β40 isoform in whole brain extracts. Finally, serum FSH levels <8 ng/mL in 16–month–old *APP*/*PS1* mice were associated with better retrieval of spatial memory. Collectively, the data provide compelling genetic evidence for a protective effect of inhibiting FSH signaling on the progression of spatial and recognition memory deficits in mice, and lay a firm foundation for the use of an FSH–blocking agent for the early prevention of cognitive decline in postmenopausal women.

## INTRODUCTION

Alzheimer’s disease (AD) poses a major global health crisis in an increasingly aged population, constituting around 60 to 80% of dementia cases. The neuropathology typically includes the presence of amyloid β (Aβ) plaques, neurofibrillary tangles, neuronal and synaptic loss, and neuroinflammation. Marked by progressive memory loss, profound physical disability, and impaired quality of life, women constitute ∼70% of the AD population^1^, and compared with men, have a higher life–time risk^2^, ∼3–fold higher progression rate^3^, and a broader spectrum of dementia–related symptoms^4^. However, mechanism(s) underpinning the preponderance of AD in women remain unclear. Post–menopausal reductions in estrogen have been considered causal, but there is evidence that estrogen upregulates, rather than suppresses, certain AD genes, such as *APOE4*^5^. Furthermore, depending upon the estrogenic compound used in a series of clinical trials, the data with hormone replacement have been mixed with improvement, no change, or even worsening of cognition^6–9^.

In contrast, there is increasing evidence that high post–menopausal levels of gonadotropins, notably FSH, are associated with established AD^10–13^. More importantly, certain neuropathologic features, including neuritic plaques, neurofibrillary tangles, and chronic gliosis, often begin prior to the last menstrual period––namely during the menopausal transition (ages 45 to 50 years)^14^. During this period, when a steady rise of serum FSH coincides with unperturbed estrogen levels, women show a sharp decline in memory function and increased risk of mild cognitive impairment (MCI) and dementia^15–17^. This phase also coincides with bone loss, obesity, dysregulated energy balance, and reduced physical activity^18–22^.

We and others have shown that FSH directly causes bone loss and increases body fat^23–27^. Prompted by these data, we asked the question: does FSH contribute to AD––and, if so, do the sharp, up to 10–fold increases in serum FSH across and beyond the menopausal transition account for the disproportionately high incidence of AD in aging women *versus* men, who display only a 3.5% annual rise in serum FSH^28^? Using AD–prone *3xTg* mice, we found that recombinant FSH or ovariectomy (high serum FSH) induce Aβ and phosphorylated TAU (pTAU), inflammation, neuronal apoptosis, and spatial and recognition memory loss^29^. Downregulating the hippocampal *Fshr* inhibits ovariectomy–induced AD pathology^29^. To block FSH action, we designed and generated a panel of polyclonal and monoclonal antibodies to a small FSHR– binding epitope of FSHβ that block FSH action^25, 30–32^. Injected into *3xTg* mice, our polyclonal antibody prevented ovariectomy–induced AD–like pathology and spatial memory loss^29^––thus, providing further, more compelling evidence that FSH is a disease driver for AD. More recently, we found that FSH interacts with the *Apoe4* gene, and not the *Apoe3* gene, in mice to promote AD–like features^33^. It also stimulates the transcription factor C/EBPβ to upregulate asparagine endopeptidase (AEP) that acts as a δ-secretase to cleave amyloid precursor protein (APP) and TAU^29, 33^.

Here, we explore whether the global deletion of the *Fshr*, which we find is expressed predominantly in AD–vulnerable brain regions^29, 34^, namely the granular layer of the hippocampal dentate gyrus and the entorhinal cortex, can prevent the onset and severity of the memory impairment in AD–prone mice. For this, we generated *Fshr* haploinsufficient and null mice on a *3xTg* background. Using the Morris Water Maze test, we documented an impressive gene–dose–dependent prevention of both the acquisition and retrieval of spatial memory loss. However, in 10–month–old ovariectomized *3xTg;Fshr^-/-^*mice, prevention was restricted to the retrieval of consolidated spatial and recognition memory. In a second experimental prong, we used 15–month–old *APP*/*PS1* mice to demonstrate a clear effect of low serum FSH levels (<8 ng/mL) in improving the retrieval of spatial memory. Given that we now have an FSH–blocking antibody that prevents memory loss in *3xTg* mice, these genetic prevention data provide a firm framework for testing our humanized monoclonal antibody, MS-Hu6^30^, for the prevention of AD and MCI in people.

## RESULTS

Here, we report, in loss–of–function studies, that the genetic deletion of *Fshr* globally in mice results in a gene–dose–dependent prevention of impairments in spatial and recognition memory. For this, we first crossed *Fshr*^+/-^ mice with *3xTg* mice to obtain *3xTg;Fshr^+/-^* mice, which were then crossed to generate *3xTg;Fshr^+/+^, 3xTg;Fshr^+/-^*and *3xTg;Fshr^-/-^* mice. Of note is that the *3xTg* background is homozygous for four mutations, namely *APP^K^*^670^*^N, M^*^671^*^L^*, *MAPT^P^*^301^*^L^* and *Psen1^M^*^146^*^V^*. As female *Fshr*^-*/*-^ mice are known to be hypogonadal, we implanted 90–day, slow–release 17β–estradiol pellets into 8–week–old mice; this, we have found, normalizes serum estradiol levels^29^. We also confirmed a near–complete loss and an ∼50% reduction of *Fshr* in *3xTg;Fshr^-/-^* and *3xTg;Fshr^+/-^*mice, respectively, on digital PCR (Fig. S1A).

We first studied the effects of depleting *Fshr* on the *3xTg* background on spatial acquisition and memory using the Morris Water Maze Test. 5–month–old female *3xTg;Fshr^+/+^*mice were expectedly impaired during the spatial acquisition phase (platform submerged, Fig. 1A), indicative of an impaired spatial learning, as well as during the retention phase (platform removed, Fig. 1B), indicating impaired spatial memory retrieval. The complete loss of the *Fshr* in *3xTg;Fshr^-/-^* mice resulted in a remarkable reduction of latency to locate the platform in the acquisition phase, statistically significant at days 3, 4 and 5, as well as in the retention phase. Partial depletion of the *Fshr* in *3xTg;Fshr^+/-^* mice also resulted in prevention of impaired learning at day 5, and a trend in the retention phase. No effect on motor activity (swim speed) was noted for any experimental group in both spatial acquisition and retention tests (Fig. S1B). Together, the data demonstrate unequivocally an effect of FSHR signaling on both spatial acquisition and memory retrieval, further substantiating the effects of pharmacologic inhibition by our FSH– blocking antibody^29^.

**Figure 1:**
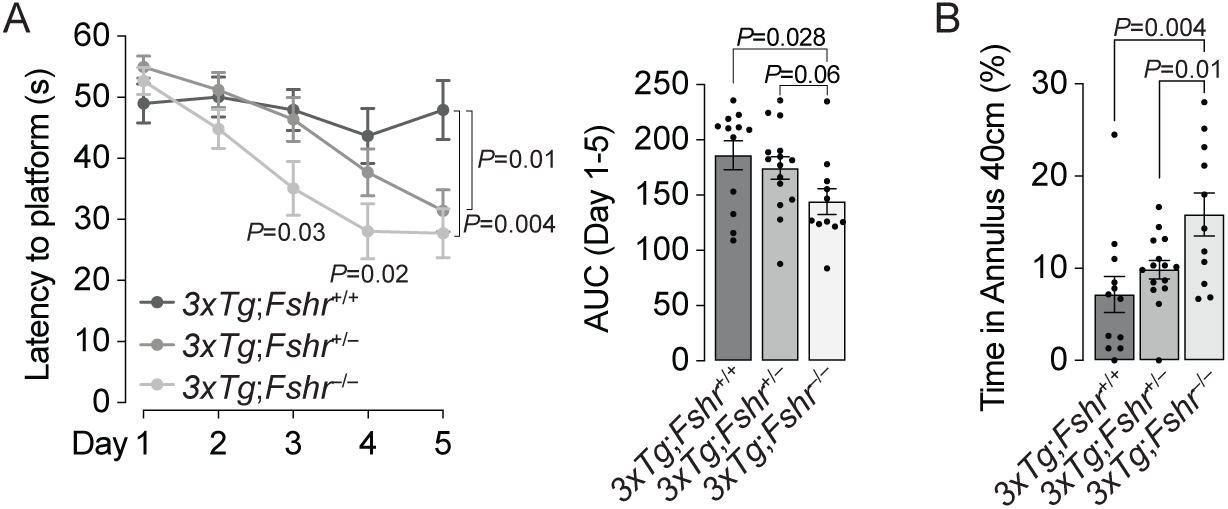
Morris Water Maze to evaluate the effect of genetic *Fshr* depletion in 5–month–old *3xTg* females on acquisition and retrieval of spatial memory. (**A**) In 5–month–old *3xTg* female mice, latency to find the hidden platform (seconds, s) was significantly shorter for *3xTg;Fshr^-/-^*on days 3, 4 and 5 of the acquisition trials compared with *3xTg;Fshr^+/+^*mice. The effect was also significant on day 5 in *3xTg;Fshr^+/-^* mice compared with *3xTg;Fshr^+/+^*mice. Both time courses and area under the curve (AUC, days 1 to 5) are shown. Mice that showed repeated episodes of extensive floating (>10 s *per* trial or >25% of trial time across 5 days) were excluded from the analysis (**B**) The effect of *Fshr* depletion in the same mouse groups at day 7 in the retention trial which platform was removed and time spent around the 40–cm platform center point (platform zone) was determined. Mice that showed repeated episodes of extensive floating (>25% of trial time) were excluded from the analysis. *N*=8 to 13 mice *per* group; Mean ± SEM; *P* values as shown, significant at *P*≤0.05; repeated measures ANOVA followed by Fisher’s Least Significant Difference post–hoc analyses).

We next evaluated the effect of *Fshr* depletion on the progression of spatial acquisition impairment with age in mice that had been ovariectomized or sham–operated at 8 weeks of age. Sham–operated 10–month–old *3xTg;Fshr^+/+^* mice expectedly displayed greater latency than at 5 months. Furthermore, 5– and 10–month–old sham–operated *3xTg;Fshr^-/-^* mice showed reduced latency compared with *3xTg;Fshr^+/+^* mice on training days 3 to 5 (Fig. 2A) and on days 4 and 5 (Fig. 2B), respectively. This indicates that *Fshr* deletion in sham–operated mice ameliorates the spatial acquisition impairment, as in Fig. 1, and its progression with age. While sham–operated heterozygotic *3xTg;Fshr^+/-^* mice showed no effect on the latency at 5 months (Fig. 2A), at 10 months, there was a significantly shorter escape latency on training day 5 (Fig 2B). In contrast, and surprisingly, ovariectomy masked the effect of *Fshr* gene depletion, resulting in no latency differences between groups during all training days at 5 months of age (Fig. 2C). However, at 10 months, there was a significantly shorter latency in ovariectomized *3xTg;Fsh^+/-^*mice at day 4, and a trend toward significance trend toward significance on days 3 and 5 compared with *3xTg;Fshr^+/+^* mice (Fig. 2D). No effect on motor activity (swim speed) was noted during any training day for all experimental groups (Fig. S1C).

**Figure 2:**
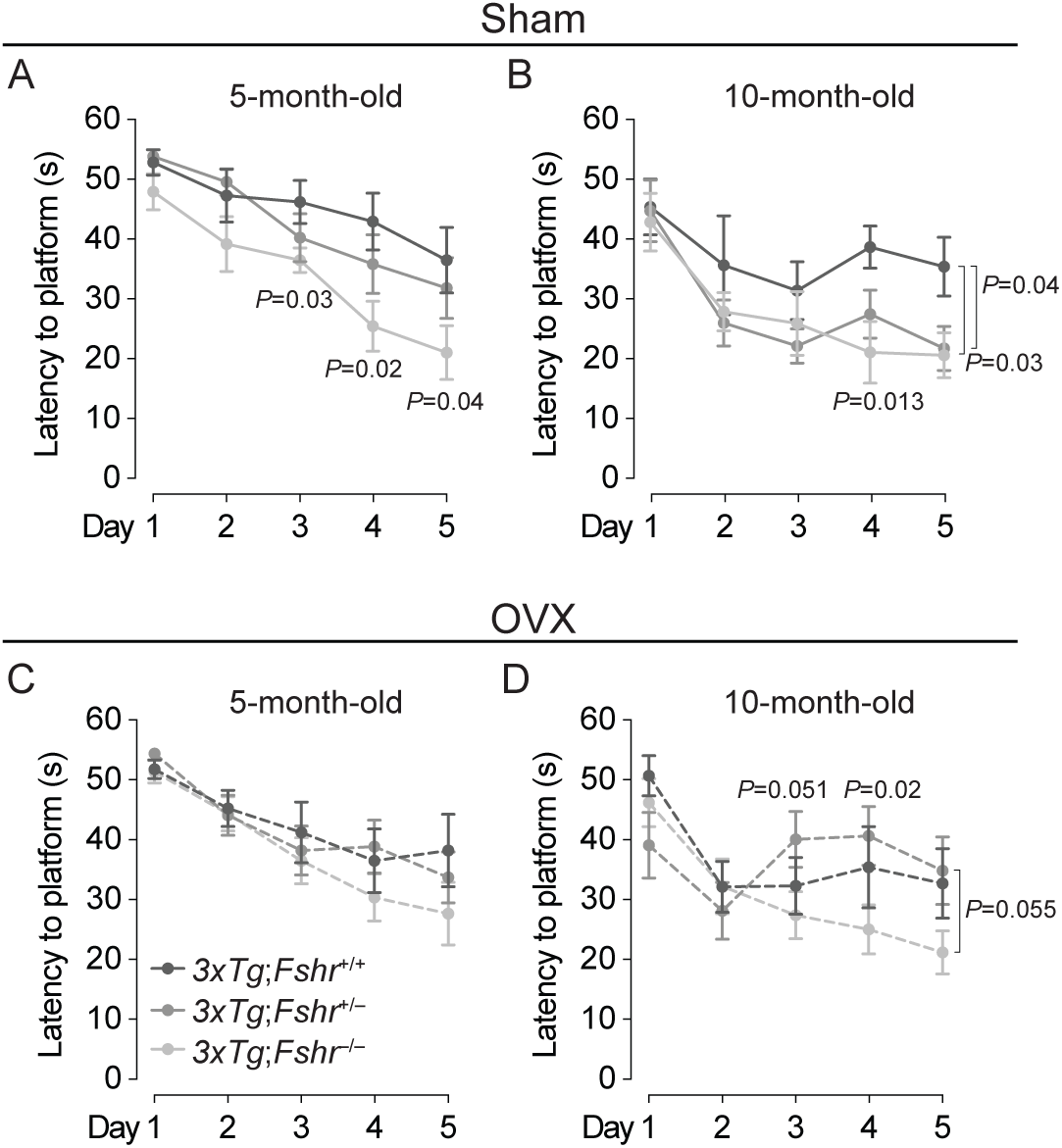
Morris Water Maze to evaluate the effect of age, ovariectomy, and genetic *Fshr* depletion in *3xTg* females on acquisition of spatial memory. In 5–month–old sham–operated *3xTg* mice (**A**), latency to find the hidden platform (seconds, s) was significantly shorter for *3xTg;Fshr^-/-^*on days 3, 4 and 5 of the acquisition trials compared with *3xTg;Fshr^+/+^*mice. At 10 months (**B**), the effect was significant on days 4 and 5 in *3xTg;Fshr^-/-^* mice and on day 5 in *3xTg;Fshr^+/-^* mice. In ovariectomized mice, there was no difference between the groups at 5 months of age (**C**), but at 10 months (**D**), the *3xTg;Fshr^-/-^*mice showed shorter latency compared with the *3xTg; Fshr^+/-^* group at days 4 and a trend toward significance on days 3 and 5. Mice that showed repeated episodes of extensive floating (>10 s *per* trial or >25% of trial time across 5 days) were excluded from the analysis. *N*=10 to 14 mice *per* group; Mean ± SEM; *P* values as shown, significant at *P*≤0.05; repeated measures ANOVA followed by Fisher’s Least Significant Difference post–hoc analyses).

We also studied the effect of *Fshr* depletion on retrieval of consolidated memory using the retention trial of the Morris Water Maze test in the *3xTg*;*Fshr* genotypes. Memory retrieval, assessed by the percent of time spent in the platform zone, was more impaired at 10 months of age compared with 5 months in both sham–operated and ovariectomized *3xTg;Fshr^+/+^* mice (Fig. 3). However, both 5– and 10–month–old *3xTg; Fshr^-/-^* mice displayed improved memory retrieval compared with *3xTg;Fshr^+/+^* mice; this effect was gene–dose–dependent (Figs. 3A and 3B). The data suggest a protective effect of graduated *Fshr* depletion in ameliorating the memory retrieval decline with age. Yet again, in 5–month–old ovariectomized mice, the beneficial effect of *Fshr* depletion on memory retrieval was absent (Fig. 3C); however, better memory retrieval was found at 10 months of age in ovariectomized *3xTg;Fshr^-/-^* mice compared with *3xTg;Fshr^+/-^* mice (Fig. 3D)—this suggests that absence of FSHR signaling does have an effect in attenuating the progression of memory retrieval decline with age in ovariectomized *3xTg* mice. No effect on motor activity (swim speed) was found during the memory retention trial for any experimental group at both ages (Figs. S1B and S1C, day 7)

**Figure 3:**
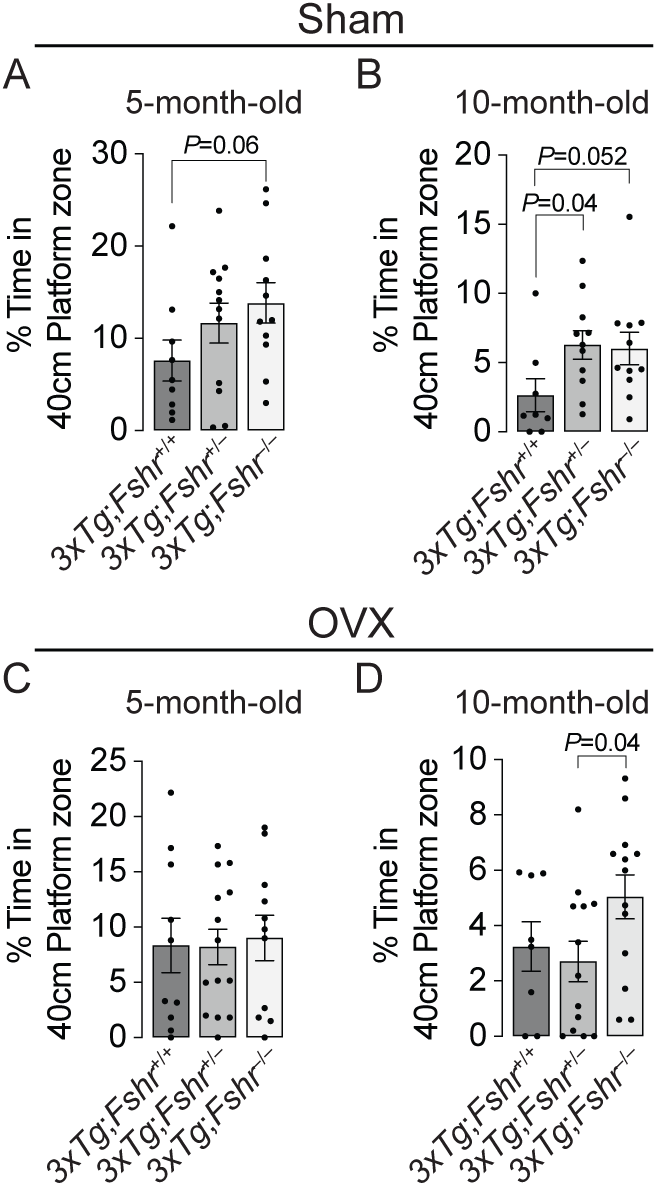
Morris Water Maze to evaluate the effect of age, ovariectomy, and genetic *Fshr* depletion in *3xTg* females on retrieval of spatial memory. In 5–month–old sham–operated *3xTg* mice (**A**), time spent in the 40–cm platform zone in the absence of the platform showed a trend to significance for *3xTg;Fshr^-/-^* mice compared with *3xTg;Fshr^+/+^* mice. At 10 months (**B**), the effect was significant for *3xTg;Fshr^+/-^*compared with *3xTg;Fshr^+/+^* mice and showed a trend toward significance in *3xTg;Fshr^-/-^* mice compared with *3xTg;Fshr^+/+^*mice. In ovariectomized mice, there was no difference between the groups at 5 months of age (**C**), but at 10 months (**D**), the *3xTg;Fshr^-/-^*mice showed longer time spent in the platform zone compared with the *3xTg;Fshr^+/-^*group. Mice that showed repeated episodes of extensive floating (>25% of trial time) were excluded from the analysis. *N*=10 to 14 mice *per* group; Mean ± SEM; *P* values as shown, significant at *P*≤0.05; repeated measures ANOVA followed by Fisher’s Least Significant Difference post–hoc analyses).

To further explore whether graduated *Fshr* loss benefited other memory domains in 10– month–old *3xTg* mice, we tested recognition memory using the Novel Object Recognition test, which is based on the inherent ability of rodents to explore and recognize a novel object in the environment over a familiar one^24^. We observed no differences between the interaction with familiar and novel objects in 10–month–old sham–operated mice of any genotype, consistent with our prior data using an FSH–blocking antibody in 9–month–old male *APP*/*PS1* mice^29^. However, significantly increased interaction with the novel over the familiar object was noted in ovariectomized *3xTg; Fshr^-/-^* group, confirming a protective effect of absent FSHR signaling on recognition memory in ovariectomized aged mice (Fig. 4). The latter finding is also consistent with the prevention of consolidated spatial memory retrieval in these mice (Fig. 3D).

**Figure 4:**
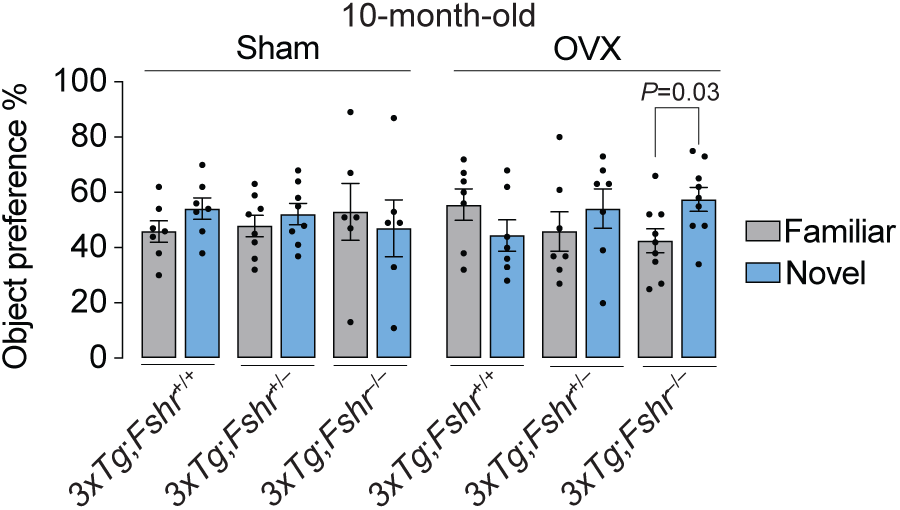
The effect of *Fshr* depletion in 10–month–old *3xTg* females following sham operation (Sham) or ovariectomy (OVX) on recognition memory in the Novel Object Recognition Test. Significant preference of a novel (N) object over a familiar (F) object represents intact recognition memory. While recognition memory is expectedly impaired in all groups at 10 months of age, ovariectomized *3xTg;Fshr^-/-^* mice showed a significant preference towards novel object, suggesting intact recognition memory. Mice that showed a total object interaction of <5% during training or testing trials were excluded from the analysis of the entire experiment. Sham: *N*=6-8/group, OVX: *N*=7-9/group; Mean ± SEM; \**P*<0.05; Student’s *t*-test.

For validation^25^, we first tested whether there was a preference to which side of the box two identical objects were placed. We found no left or right preference in the identical object trial irrespective of genotype (Fig. S1D). Further validation of our dataset required that mice interact with both objects at a minimal threshold of 5% during both training and testing trials; mice that explored below this threshold were excluded *a priori*. Object interactions ranged from 25.3% to 37.8% in the training trial and from 9.2% to 14.4% in the testing trial. There was no main effect of genotype or operation in the training and testing trials (Table S1). The observed increase in novel over familiar object interaction in ovariectomized *3xTg;Fshr^-/-^*mice is also unlikely to be due to increased general locomotor activity, since ovariectomy *per se* reduced general activity during training trials, and the testing trial had no effect on locomotion (Table S1).

We measured Aβ40 and Aβ42 isoforms in whole brain extracts using ELISA in 10– month old mice. There was a clear *Fshr* gene–dose–dependent reduction of Aβ40 levels in sham–operated and ovariectomized *3xTg* mice (Fig. 5), consistent with the prevention of memory retrieval deficit (Fig 3D). However, only sham–operated *3xTg;Fshr^-/-^* mice showed a significant reduction in Aβ42 compared with *3xTg;Fshr^+/+^* mice.

**Figure 5:**
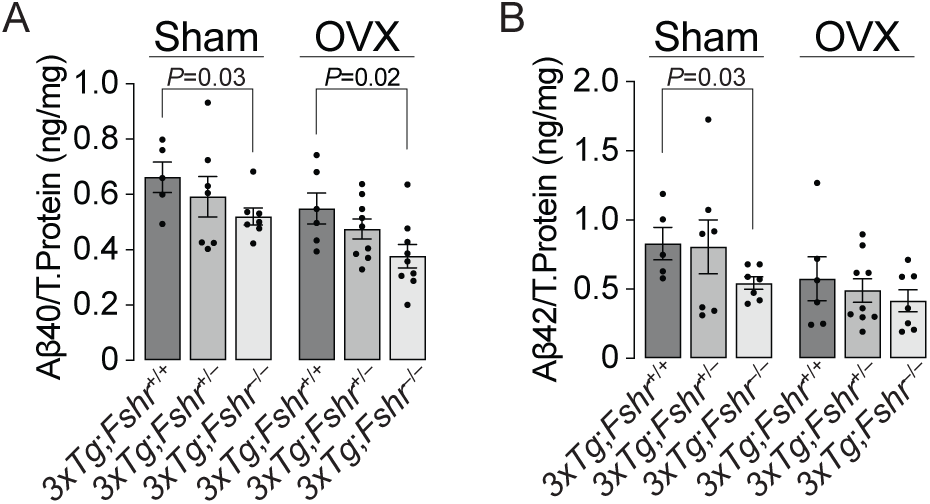
The effect of *Fshr* depletion in 10–month–old *3xTg* females following sham operation (Sham) or ovariectomy (OVX) on the accumulation of amyloid β (β) isoforms, Aβ40 (**A**) and Aβ42 (**B**) in whole brain extracts. There was a gene–dose–dependent reduction of Aβ40 accumulation in both sham–operated and ovariectomized groups, but reductions in Aβ42 were significant only in the sham–operated *3xTg;Fshr^-/-^* group. Sham: *N*=6-8/group, OVX: *N*=7-9/group; Mean ± SEM; *P* values as shown, significant at *P*≤0.05, Student’s *t*-test.

Lastly, to complement the effects of deleting the *Fshr* genetically in *3xTg* mice, we studied the effect of circulating FSH levels on acquisition and retrieval of spatial memory in 16– month–old *APP*/*PS1* mice––which, unlike *3xTg* mice, represent a less aggressive, single pathology model involving Aβ accumulation. These mice develop spatial memory impairments around 15 months^35, 36^. For this, we pooled two cohorts of *APP*/*PS1* mice and separated the combined group by an arbitrary cut–off of serum FSH at 8 ng/mL to reflect broadly pre– and post–menopausal FSH levels (Fig. S1E). The learning trials of the Morris Water Maze test revealed no difference in spatial acquisition between the low FSH (<8 ng/mL) and high FSH (>8 ng/mL) group (Fig. 6A). However, using the retention trial, we found an impressive difference between the two groups in the time spent in platform zone favoring the low–FSH group (Fig. 6B). These data provide additional evidence that low circulating FSH levels in mice are directly associated with better retrieval of consolidated memory.

**Figure 6:**
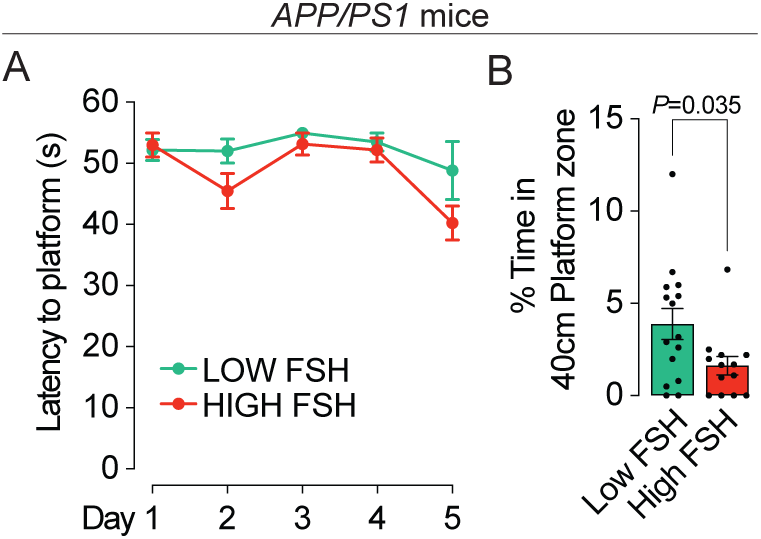
Morris Water Maze to evaluate the effect of elevated FSH in 15–month–old *APPIPS1* females on acquisition and retrieval of spatial memory. (**A**) In 16–month–old *APP/PS1* female mice, latency to find the hidden platform (seconds, s) was not different among the groups. Mice that showed repeated episodes of extensive floating (>10 s *per* trial or >25% of trial time across 5 days) were excluded from the analysis. (**B**) The effect of elevated FSH in the same mouse groups at day 7 in the retention trial which the platform was removed and time spent around the 40–cm platform center point (platform zone) was determined. Mice with low FSH levels showed significantly greater time spent in the platform zone. Mice that showed repeated episodes of extensive floating (>25% of trial time) were excluded from the analysis. *N*=13 to 15 mice *per* group; Mean ± SEM; *P* values as shown, significant at *P*≤0.05; repeated measures ANOVA followed by Fisher’s Least Significant Difference post–hoc analyses).

## DISCUSSION

There is little information on circuitry through which glycoprotein hormones from the anterior pituitary regulate central functions^37^. We recently discovered that FSH, *hitherto* considered solely a fertility hormone but with a number of newly–discovered somatic functions^25–27^, acts on FSH receptors on neurons^29^. Furthermore, our comprehensive analysis through RNAscope at the single transcript level revealed *Fshr* transcripts in 353 regions of the brain, *albeit* without ascribed functions^34^. High *Fshr* expression was noted selectively on neurons in AD–vulnerable regions, namely on the granular layer of the dentate gyrus of the hippocampus and the entorhinal cortex^29, 34^. This allowed us to interrogate the hippocampal *Fshr* through siRNA knockdown, which revealed a clear attenuation of both spatial memory acquisition and retrieval in ovariectomized *3xTg* mice^29^. Furthermore, FSH injections caused spatial memory impairment, whereas our FSH–blocking antibody attenuated the loss of both acquisition and retrieval of memory^29^. Here, we provide intriguing genetic data that unequivocally establish a role for FSH in regulating spatial memory impairment in models of AD mouse model. We report that the global loss of *Fshr* expression produces a gene–dose–dependent amelioration of defects in both the acquisition and retrieval of memory in *3xTg* mice, and that, equally importantly, *APP*/*PS1* mice with serum FSH levels ≤8 ng/mL display improved retrieval of memory.

We find an expected impairment of acquisition and retrieval of spatial memory with age in unoperated or sham–operated *3xTg* mice. Furthermore, *Fshr* depletion on a *3xTg* background causes a gene–dose–dependent attenuation of both spatial acquisition and retrieval of memory at 5 and 10 months of age. Consistent with this, Aβ40 and Aβ42 were reduced in *Fshr–*deficient mice in a gene–dose–dependent manner. This provides compelling evidence that the age–associated impairment of the two components of spatial memory––acquisition and retrieval––as well as Aβ isoform accumulation are rescued upon *Fshr* depletion.

However, in contrast to unoperated or sham–operated mice, the effect of age on memory impairment in ovariectomized was surprisingly restricted to memory retrieval, but not to the acquisition of spatial memory. Thus, we found that while memory retrieval was impaired with age, there was no difference between sham–operated and ovariectomized groups, likely due to a ceiling effect at 10 months. However, the impaired memory retrieval at 10 months showed improvement upon complete deletion of the *Fshr*, when compared with *Fshr* haploinsufficiency. This was consistent with a significant reduction in Aβ40 in *Fshr*–depleted ovariectomized *3xTg* mice. These findings are further concordant with the rescue of recognition memory––another form of hippocampus–dependent consolidated memory––in 10–month–old, *Fshr*–null mice. The results also support data showing that low *basal* serum FSH in a different mouse model, the *APP*/*PS1* mouse, is associated with better consolidated memory retrieval, but not with improved acquisition of spatial memory.

In addition to FSH, rising post–menopausal levels of LH have also been implicated in the pathogenesis of memory deficits^38^. Earlier reports show that LHβ transgenic mice or mice receiving human chorionic gonadotropin (hCG) are cognitively impaired^39, 40^. Furthermore, similarly to the *Fshr*, *Lhcgr* transcripts are expressed in AD–vulnerable regions, such as the dentate gyrus of the hippocampus and entorhinal cortex^34^. Although we recently discovered that absent LH signaling in *Lhcgr*^-/-^ mice prevents the anxiety phenotype that develops with aging, further loss–of–function studies using cognitively impaired mice should shed light on the potential effects of LH on memory functions.

With that said, our findings provide a unique genetics–based framework for FSH inhibition to prevent AD–like features in people, particularly in women across the menopausal transition where deficits in memory and MCI are associated with rapid bone loss and the onset of visceral obesity^15, 18, 19^. Towards targeting a rising FSH in these women, as well as post– menopausal women of advanced ages, we recently produced a first–in–class humanized FSH– blocking antibody, MS-Hu6, which targets a short FSHR–binding epitope of FSHβ, and, in doing so, blocks FSH action^30^. We have shown that MS-Hu6 has an acceptable affinity to FSH, with a K_D_ of 7.2 nM that approaches trastuzumab; a long half–life of 7.8 days in humanized *Tg32* mice; limited, but measurable accumulation in the brain upon subcutaneous injection; and thermal, colloidal, monomeric, structural and accelerated stability in formulation, as evidence of durability and manufacturability^30, 41, 42^. However, supporting our core concept of inhibiting FSH to prevent MCI is a recent study in women between ages 40 to 65, of which 35% were peri–menopausal––documenting a strong positive correlation between serum FSH levels and Aβ load (measured on PET scans) and gray matter volume in AD–vulnerable regions^43^.

## ACKNOWLEDGEMENTS

Work at Icahn School of Medicine at Mount Sinai carried at the Center for Translational Medicine and Pharmacology was supported by R01 AG071870, R01 AG074092 and U01 AG073148 to T.Y. and M.Z.; and U19 AG060917 and R01 DK113627 to M.Z.

## LEGENDS TO SUPPLEMENTARY FIGURES, TABLE AND VIDEOS

**<u>Figure S1</u>: (A)** *Fshr* mRNA levels were quantified using droplet digital PCR in the ovary in the various *3xTg* mutants. There was a 40-50% reduction in *Fshr* mRNA in *3xTg;Fshr^+/-^* mice and no expression in *3xTg;Fshr^-/-^* mice in compare with *3xTg;Fshr^+/+^*. *N*=5 to 8 mice *per* group; Mean ± SEM; *P* values as shown, significant at *P*≤0.05; repeated measures ANOVA followed by Fisher’s Least Significant Difference post–hoc analyses). No swim speed differences were noted across the groups of 5–month–old unoperated (**B**) or 5– or 10–month–old sham–operated or ovariectomized *3xTg*;*Fshr* mutants (**C**) on days 1 to 5 (spatial acquisition trial) or day 7 (retention trial) on the Morris Water Maze test. Mice that showed repeated episodes of extensive floating during the acquisition (>10 s *per* trial or >25% of trial time across 5 days) or retention trial (>25% of trial time) were excluded *a priori* from the analysis. *N per* Figs. 1 to 3. Mean ± SEM; repeated measures ANOVA followed by Fisher’s Least Significant Difference post hoc analyses. **(D)** No side preferences were found across all groups of 10–month–old *3xTg*;*Fshr* mutants on a training trial conducted with identical objects 6 hours before the Novel Object Recognition test. Mice that showed total object interaction of less than 5% during training or testing trials were excluded *a priori* from analysis of the entire experiment. *N per* Fig. 4. Mean ± SEM; Student’s unpaired *t*-test. **(E)** Serum FSH levels in 16–month–old female *APP*/*PS1* mice using an ELISA. Animals were pooled from two cohorts using a cut–off of 8 ng/mL.

**<u>Table S1</u>:** Statistical analyses for the Novel Object Recognition test for 10–month–old sham– operated or ovariectomized *3xTg*;*Fshr* mutants. Noted are total distances traveled and percent of object interactions in the training and testing phases of the test. Mean ± SEM; repeated measures ANOVA followed by Fisher’s Least Significant Difference post–hoc analyses.

**<u>Supplementary Video 1</u>:** Video showing an unimpaired *3xTg* mouse reaching the hidden platform within 10 seconds in the Morris Water Maze test.

**<u>Supplementary Video 2</u>:** Video showing an impaired *3xTg* mouse that was unable to reach the hidden platform during the testing duration of 60 seconds in the Morris Water Maze test.

## METHODS

### Mouse Models

*3xTg* mice were sourced from Jackson Laboratory (strain: 034830). The mice carry a transgene containing mutated human *APP*^K670N/M671L^ and *MAPT*^P301L^, as well as a knock–in mutation *Psen1*^M146V^, on a heterozygous C57BL*/6*;129 background^44^. The mice exhibit AD–like neuropathology and a decline in long–term memory around 3 to 4 months of age^26^. *Fshr* mutants were bred and maintained at the Icahn School of Medicine at Mount Sinai (ISMMS), with heterozygotes on 129T2svEmsJ^27^. The two strains were crossed to produce viable F1 hybrid *3xTg*^+/-^*;Fshr*^+/-^ littermates. The latter were subsequently crossed with *3xTg*^+/+^ mice to generate compound *3xTg*^+/+^*;Fshr^+/-^*mice, which were then crossed to create *3xTg*^+/+^*;Fshr*^+/+^; *3xTg*^+/+^*; Fshr^+/-^* and *3xTg*^+/+^*;Fshr^-/-^* mice (*hitherto* termed *3xTg;Fshr^+/+^*, *3xTg;Fshr^+/-^*, and *3xTg;Fshr^-/-^* mice). Half of the animals in each group underwent either ovariectomy or sham operation. 90–day, slow–release pellets containing 0.36 mg 17β-estradiol were inserted in the unoperated and sham–operated *3xTg;Fshr^-/-^* to normalize their estrogen level, as before^29^. *APP/PS1* mice were obtained from Jackson Laboratory (strain: 34829) and maintained at ISMMS. The mice carry a human transgene containing *APP*^K670N/M671L^ and *PSEN^1^*^Δ*E*9^ mutations^36^. All experimental mice were grouped–housed to reduce single–house stress, under a 12–hour light/dark cycle with food and water *ad libitum*. Behavioral tests were performed at ages 5 and 10 months for the *Fshr*;*3xTg* mutants, and at 15 months for *APP*/*PS1* mice. All tests were conducted in the light phase, and the order of behavioral tests was the same for each mouse. The protocols were reviewed and approved by the ISMMS Institutional Animal Care and Use Committee.

### Surgery

For ovariectomy and sham operation, mice were anesthetized using ketamine/xylazine and received a prophylactic dose of meloxicam to alleviate potential post-surgical pain or distress. After anesthesia, the lower back was shaved, cleaned with 70% ethanol, and washed with povidone prior to the surgery. A ∼1 cm skin incision was made, followed by an incision through the muscle layer to access the peritoneal cavity. The ovaries were identified, extracted one at a time through the incision, tied off, and removed. The muscle layer was sutured, and the external incision was closed using wound clips. For pellet implantation, a 0.5 cm incision was created in the skin at the nape of the neck of the *3xTg;Fshr^-/-^*mice. A small pocket was carefully dissected towards the caudolateral area behind the ear, where the 90–day, slow– release pellet containing 0.36 mg 17β-estradiol was placed using tweezers. The incision was closed with a wound clip. Each mouse was placed in a clean cage, allowed to recover from anesthesia, and returned to the home cage after exhibiting normal behavior and ambulation. Meloxicam was continued 24, 48, and 72 hours after surgery.

### Reagents

ELISA kits for human Aβ40 (Cat. #KHB3481) and Aβ42 (Cat. #KHB3544) were purchased from Invitrogen and FSH (Cat. #MPTMAG-49K) were purchased from Millipore. The 90–day, slow–release pellets containing 0.36 mg 17β-estradiol were purchased from Innovative Research of America (Cat. #NE121).

### Digital PCR

*Fshr* mRNA levels were quantified using droplet digital PCR (ddPCR). In brief, RNA was isolated by TRIzol (Life Technologies). Reverse transcription was performed using SuperScript III reverse transcriptase (Life Technologies). Isolated RNA was used to perform ddPCR to determine *Fshr* mRNA levels using FSHR Probe (Life Technologies, Cat. #4331182). Droplets containing the cDNA were generated using a Biorad Droplet generator (QX200) by mixing with droplet generator oil (Cat. #D9161172A), and the formed droplets were amplified using a thermocycler (Applied Biosystems) and analyzed using droplet reader (Biorad).

### Behavioral Tests

Behavioral testing, described by us previously^45^, consisted of two memory tests conducted in the order of increasing invasiveness in the following order: Novel Object Recognition and Morris Water Maze. Mice received 3 days of resting time between tests to decrease carryover effects from prior tests. Mice were habituated to the testing room for 30 minutes at the beginning of each test day. The order of tests in which mice were tested was the same across all mice. Each mouse was tested once *per* test. The behavioral room wall cues remained the same for all tasks and the same experimenter conducted all of the tests. All test trials were video–recorded, tracked, and analyzed with ANY-maze tracking software (v 7.2; Stoelting, Wood Dale, IL). Locomotor activity data for each test are summarized in Table S1.

### Novel Object Recognition Test

The Novel Object Recognition test was performed in square test boxes (40×40×35 cm) with even lighting conditions (30 ± 5 lux). Each test box consisted of grey steel bottom plate, white Perspex under a camera mounted above all boxes. A tower of Lego bricks and Falcon tissue culture flask filled with sand were used as objects^27^. Prior to the experiments, both objects were tested with a separate cohort of mice to exclude that mice showed a preference for either object or side preference due to the behavioral room conditions. Sample object and the novel object placement followed a counterbalanced design between trials to control for order and location effects.

The test consisted of two trials––training and testing––separated by 6 hours. In the training trial, mice were placed into the test box containing two equal sample objects (e.g., flasks), in front of the south wall facing away from the objects. Each mouse was allowed to explore the objects for 10 minutes before it was returned to its home cage. After 6 hours, the testing was conducted by placing the mouse into the same test box again, but containing one sample (familiar) object and one unfamiliar object (a flask and a Lego bricks tower) and object interaction was recorded for 10 minutes. After each trial, the objects and boxes were cleaned with a Quatricide dilution to eliminate odor cues. The maze was cleaned using 70% ethanol between each trial. Object interaction was defined as an event where a mouse’s head was within 2 cm of the object and directed towards the object, excluding sitting on the objects^28,29^. For the training trial, object Interaction [%] was calculated as [sample object interaction time]/[total test time] x 100%. For the testing trial, object interaction [%] was calculated as [novel object interaction time]/[total object interaction time] x 100%^30^. Mice with less than 5% of total object interaction in either trial were excluded from the analysis^31^.

### Morris Water Maze

To test spatial memory accusation and retrieval, we used the Morris Water Maze test (adapted from Vorhees *et al.*)^32^. This utilized a circular pool (150 cm diameter) filled with water (26 ± 1°C; 10 cm distance from water surface to wall rim) made opaque with non–toxic tempera paint. A circular rescue platform (diameter: 11 cm; distance between platform center point and pool wall: 27 cm) was submerged 1–1.5 cm below the water surface and the testing area was illuminated with indirect lighting (150 ± 10 lux) to avoid reflections. To monitor animals during trials, a camera was mounted to the ceiling centrally above the pool. The water maze was surrounded by black–and–white extra–maze cues on the walls of the room. Repeated episodes of excessive floating (>10 seconds and/or ≥25% of trial across five days of training) was rare and found in only 6 mice during the entire study. These mice were excluded from the analysis *a priori*^32^.

For spatial acquisition trials, a submerged rescue platform, invisible to the mice was used. To locate the escape platform, mice used the extra–maze cues. The platform location remained the same for all trials, whereas the starting location was varied between trials. Mice had 60 seconds to find the rescue platform, after which they were guided there. Each mouse performed four trials *per* day over 5 days with an inter–trial interval of 15 to 20 minutes. Mice that failed to locate the platform during the 60 second trial, were placed on the platform for 15 seconds immediately after the end of the trial. At the end spatial acquisition day 5, mice were housed back in home cage for 24 hours (day 6). The retention trial was conducted on day 7 without additional training. For the retrieval trials, the rescue platform was removed from the pool and the mouse was allowed to swim for 60 seconds. Mice with extensive floating of >25% of the trial time were removed from the entire analysis *a priori*. For the spatial acquisition trials, the mean latency to reach the platform was calculated for each test. For the retention trials, the percent of the time spent in the platform zone (40 cm diameter surrounding the platform center point) was analyzed^32^ (See Supplementary Videos 1 and 2 for examples of unimpaired and impaired *3xTg* mice in the Morris Water Maze testing).

### Statistical Analysis

Statistical analyses were performed using GraphPad Prism v.10. For molecular analyses, the tests were either unpaired two-tailed Student’s *t*-test (two-group comparison) or one–way ANOVA followed by Fisher’s least significant difference post hoc test (more than two groups). Differences with *P*_≤_0.05 were considered significant. *P* values are annotated in the figures and are provided in the Source Data Files. For behavioral analyses, repeated measures two–way ANOVA followed by Fisher’s Least Significant Difference post–hoc test (more than two groups), one–way ANOVA or two–tailed Student’s *t*-test (two–group comparison) were utilized. Differences with *P* ≤0.05 were considered significant.

