## Supplementary figures and images for "Gene–Dose–Dependent Reduction *Fshr* Expression Improves Spatial Memory Deficits in Alzheimer’s Mice"

### Supplementary Figure 1

#### Supplementary Figure 1

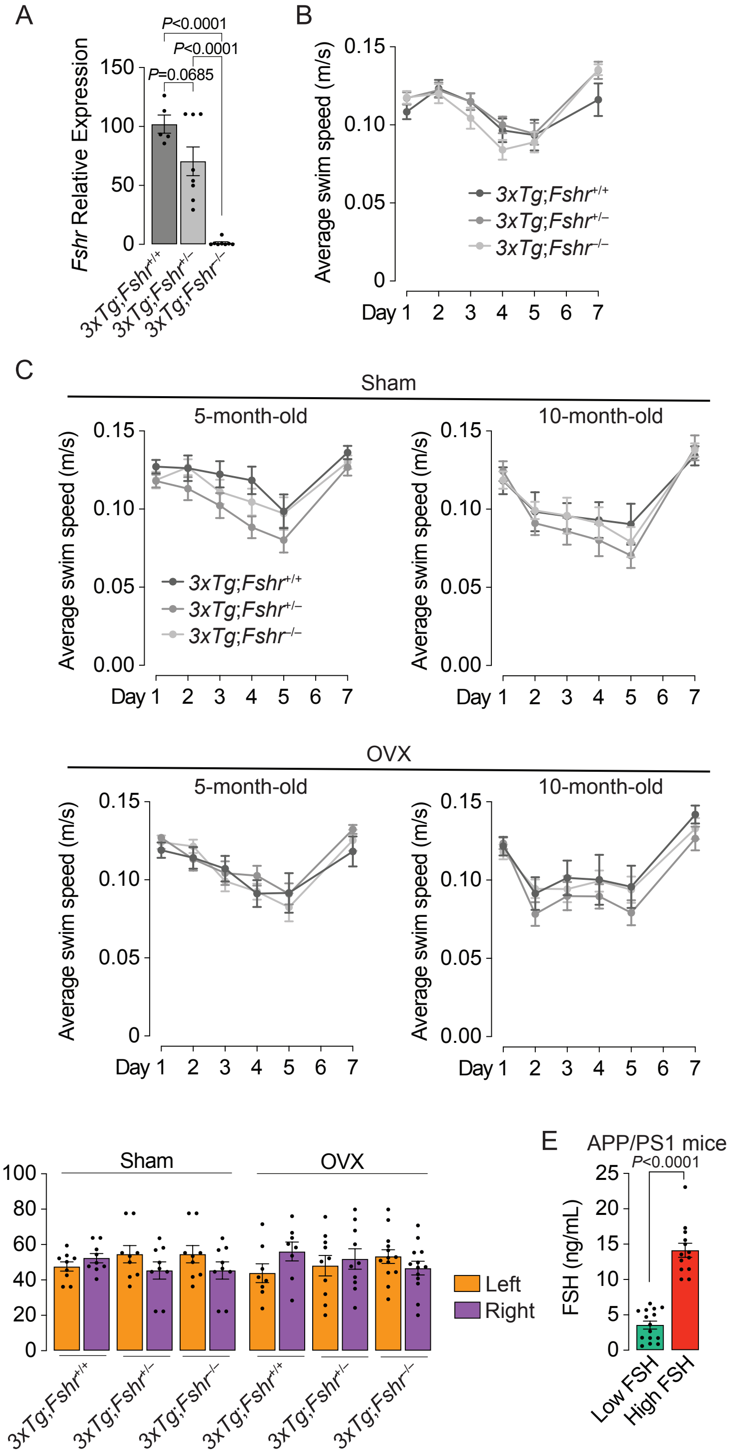
