## Supplementary material for "Gene–Dose–Dependent Reduction *Fshr* Expression Improves Spatial Memory Deficits in Alzheimer’s Mice": Table S1

| | Sham | | | OVX | | | 2- WAY ANOVA | F (DFn, DFd) | *P* value |
|---|---|---|---|---|---|---|---|---|---|
| | ***Fshr***$^{+/+}$ | ***Fshr***$^{+/–}$ | ***Fshr***$^{–/–}$ | ***Fshr***$^{+/+}$ | ***Fshr***$^{+/–}$ | ***Fshr***$^{–/–}$ | | | |
| Total distance traveled in Training Trial (m) | | | | | | | | | |
| ***N*** | 9 | 9 | 9 | 8 | 11 | 12 | **Interaction** | F (2, 52) = 2.766 | *P*=0.072 |
| **Mean** | 37.77 | 29.94 | 26.49 | 25.28 | 27.48 | 24.46 | **Operation** | F (1, 52) = 8.032 | *P*=0.007 |
| **SEM** | 2.91 | 2.54 | 2.35 | 1.24 | 2.68 | 2.13 | **Genotype** | F (2, 52) = 3.027 | *P*=0.057 |
| Percent of objects' interaction in Training Trial | | | | | | | | | |
| ***N*** | 9 | 9 | 9 | 8 | 11 | 12 | **Interaction** | F (2, 52) = 0.008 | *P*=0.992 |
| **Mean** | 15.77 | 14.55 | 12.33 | 13.44 | 11.98 | 10.18 | **Operation** | F (1, 52) = 2.830 | *P*=0.099 |
| **SEM** | 2.16 | 1.53 | 1.83 | 1.90 | 1.94 | 0.80 | **Genotype** | F (2, 52) = 1.932 | *P*=0.155 |
| Total distance traveled in Testing Trial (m) | | | | | | | | | |
| ***N*** | 7 | 6 | 8 | 7 | 9 | 7 | **Interaction** | F (2, 38) = 1.718 | *P*=0.193 |
| **Mean** | 28.14 | 23.81 | 12.63 | 18.87 | 28.61 | 16.49 | **Operation** | F (1, 38) = 0.003 | *P*=0.954 |
| **SEM** | 5.30 | 2.76 | 0.71 | 1.96 | 8.22 | 1.86 | **Genotype** | F (2, 38) = 4.214 | *P*=0.022 |
| Percent of objects' interaction in Testing Trial | | | | | | | | | |
| ***N*** | 7 | 6 | 8 | 7 | 9 | 7 | **Interaction** | F (2, 38) = 0.687 | *P*=0.509 |
| **Mean** | 14.14 | 13.13 | 13.50 | 14.14 | 15.14 | 9.22 | **Operation** | F (1, 38) = 0.425 | *P*=0.518 |
| **SEM** | 3.53 | 1.66 | 4.58 | 3.53 | 3.52 | 0.997 | **Genotype** | F (2, 38) = 0.522 | *P*=0.598 |

**Table S1:** Statistical analyses for the Novel Object Recognition test for 10–month–old sham–operated or ovariectomized *3xTg*;*Fshr* mutants.
